# Immunomodulatory Effects of Insulin-Derived Fibrils from Infusion Pumps: Role of Phenolic Preservatives in Macrophage Activation

**DOI:** 10.64898/2026.06.04.726295

**Authors:** Priscila Silva Cunegundes, Congxiao Cheng, Elizabeth Wisman, Daniel L. Menkes, Ulrike Klueh

## Abstract

**Background:** Protein fibrillation represents a critical challenge in therapeutic insulin delivery, yet the structural determinants and immunological consequences of insulin-derived fibrils (IDFs) formed in the presence of phenolic preservatives remain poorly characterized.

**Objective:** This study investigated the structural characteristics of IDFs formed with (IDF (+)) and without (IDF (-)) phenolic preservatives and elucidated their differential immunomodulatory mechanisms in bone marrow-derived macrophages (BMDMs).

**Methods:** IDF structural properties were characterized using Thioflavin T fluorescence and nanoparticle tracking analysis (Spectradyne nCS1™). BMDMs were treated with serial dilutions of IDF (+), IDF (-), or m-cresol. Cytotoxicity, reactive oxygen species (ROS) production, MIP-1α levels, and expression of signaling pathways were quantified.

**Results:** Structural analysis revealed similar aggregation states between IDF (+) and IDF (-). However, IDF (+) induced greater cytotoxicity and ROS production than IDF (-), which produced minimal ROS. Both fibrils increased MIP-1α chemokine levels. Additionally, IDF (-) upregulated NRF2 whereas m-cresol downregulated STAT6 compared to control. Together, these results support the existence of distinct mechanisms of macrophage activation and suggest that protein aggregates can directly induce macrophage responses independent of ROS production.

**Conclusions:** Insulin fibrils activate macrophage inflammatory pathways through ROS-independent mechanisms. Phenolic preservatives enhance fibril cytotoxicity and likely ROS production while differentially modulating inflammatory signaling. These findings suggest that strategies to remove or reduce the effects of IDFs in insulin infusion therapy may increase longevity and biocompatibility of these devices.

## 1 Introduction

Advances in insulin delivery technologies, (e.g., infusion pumps and automated insulin delivery systems), have improved glycemic control significantly resulting in reduced diabetic complications. However, standard infusion sets used in type 1 and insulin-dependent type 2 diabetes have a limited lifespan, primarily due to infusion set-related complications, including occlusions, kinking, catheter blockage, adhesive failure, skin irritation, and lipohypertrophy ^1^. In an international survey of 14,015 continuous subcutaneous insulin infusion (CSII) users, 42% reported site-related issues ^2^. Another study in children and adolescents documented high rates of cannula blockages (64.4%), needle dislodgment (39%), and skin reactions (54.2%) ^3^. Currently, infusion sets are FDA approved for 2–3 days of wear with steel or Teflon cannulas and up to 7 days with extended-wear designs ^4^. Nonetheless, many patients replace infusion sets earlier due to infusion-site complications, most notably deterioration in glycemic control. This short lifespan is a major constraint such that next-generation insulin infusion pump technologies need to improve reliability and reduce patient burden. Among these complications, biochemical changes to insulin itself, particularly protein aggregation, represent a critical but incompletely understood negative contributing factor to infusion set failure.

Protein aggregation, or the formation of insulin-derived fibrils (IDFs), is a known complication of pump therapy ^5,6^. IDFs form when insulin molecules self-associate into amyloid-like structures, losing their native, functional conformation. This misfolding and aggregation can occur through several mechanisms, including material interactions, liquid-air interfaces, and mechanical stressors within the infusion system ^7–9^. Fibril-related issues are associated with occlusions, pump malfunctions, and inflammatory tissue reactions ^10–14^. These aggregates impair insulin bioavailability and can adversely impact diabetes management ^15^.

Despite current mitigation strategies such as site rotation and patient education, IDF-related complications persist, such that major diabetes organizations now recognize IDFs as a barrier to prolonged infusion set use ^4,16^. While IDFs are well documented in pump systems ^6,13^, the mechanisms by which these fibrillar aggregates interact with biological tissues and initiate inflammatory responses remain incompletely understood.

Macrophages are key mediators of the inflammatory response and primary responders to foreign materials at tissue-device interfaces ^17,18^. Once activated, macrophages can trigger acute or chronic inflammatory cascades ^19^. Given the tissue reactions associated with IDF deposition at infusion sites, understanding the mechanisms by which IDFs influence macrophage function is critical for informing the rational design of infusion systems. An additional consideration in IDF formation is the role of pharmaceutical preservatives. Commercial insulin formulations are manufactured in bulk and require preservatives to ensure extended shelf life and sterility. All current insulin formulations contain phenolic compounds as preservatives, (e.g. m-cresol) ^20–22^. However, these preservatives may adversely influence the biological response to aggregated insulin. To investigate these effects, BMDMs serve as a primary screening platform for understanding IDF’s effects on innate immune cells. In this study, inflammatory gene expression, intracellular signaling pathways, and particle size distributions were analyzed systematically to compare macrophage responses to IDFs formed in the presence (IDF (+)) or absence (IDF (-)) of phenolic preservatives. The goals were to elucidate the relationship between fibril structural characteristics and macrophage activation, assess preservative-dependent effects on IDF immunogenicity, and determine cytotoxicity and immunomodulatory responses. Collectively, these findings will provide mechanistic insight into IDF-mediated inflammation. Addressing these mechanisms will optimize exogenous insulin administration thus improving the clinical efficacy of insulin pump therapy.

## 2 Methodology

### 2.1 Experimental Animals and Ethical Approval

Male C57BL/6 mice aged 8-12 weeks were used as bone marrow donors for macrophage isolation. All animal procedures were conducted in accordance with the Guide for the Care and Use of Laboratory Animals and were approved by the Institutional Animal Care and Use Committee (IACUC) at Wayne State University.

### 2.2 Preparation of Insulin-Derived Fibrils (IDF) and m-cresol solution

Two types of insulin-derived fibrils were generated using commercial Humalog® (Insulin Lispro Injection U-100, Eli Lilly, IN, US) which is free of bacterial endotoxins (Lipopolysaccharide - LPS) to distinguish the effects of the fibrillar structure from those of phenolic preservatives:

#### 2.2.1 IDF (+): Fibrils with Preservatives

IDF (+) was prepared using commercial Humalog®, which contains m-cresol (3.15 mg/mL), glycerol, sodium phosphate dibasic, and zinc oxide. Insulin was transferred into sterile 15 mL polypropylene centrifuge tubes, positioned at approximately 45°, and subjected to continuous orbital agitation at 300 rpm for 48 hours at 37°C.

#### 2.2.2 IDF (-): Fibrils without Preservatives

To generate preservative-free fibrils, Humalog was first desalted using Zeba™ Spin Desalting Columns (#89890, 7K MWCO, Thermo Fisher Scientific, MA, US) according to the manufacturer’s instructions. Columns were equilibrated with Dulbecco’s Phosphate-Buffered Saline (#14190-144, DPBS, Thermo Fisher Scientific) by three successive centrifugations at 1,000 × *g* for 2 minutes at room temperature (RT). This protocol is adapted from Woods et al. (2012) ^23^, which demonstrated efficient removal of phenolic preservatives.

Insulin (700 μL) was applied to the equilibrated column, and the column was centrifuged at 1,000 × *g* for 2 minutes to collect the preservative-free fraction. The desalted insulin was then subjected to the same fibrillization protocol as IDF (+). After fibril formation, IDF (-) was washed three times with sterile 0.9% saline (Sodium Chloride Injection, USP, IL, US) by centrifugation at 10,000 × *g* for 10 minutes at RT, then resuspended to its original volume in saline.

#### 2.2.3 Preparation of m-cresol Control Solution

A preservative control solution was created to match the composition of Humalog excipients, including 3.15 mg/mL of m-cresol (#110581000, Thermo Fisher Scientific) sodium phosphate dibasic (1.88 mg/ml) (#S5136, Sigma-Aldrich, MO, US), glycerol (16 mg/ml) (#G33-1, Fisher Scientific, ON, US), and distilled water, with the pH adjusted to 7.0–7.8. This solution was sterilized by filtration through a 0.22 μm membrane filter before use.

### 2.3 Fibril measurement by ThT

Thioflavin T (ThT) fluorescence detection was performed following established protocols, with minor optimizations for this study. ThT (#A0374817, Thermo Fisher Scientific) was prepared as a 219 µM working solution in sterile 0.9% saline immediately before use. All samples were dispensed into a black, clear-bottom 96-well fluorescence plate (#3603, Corning Costar®, ME, US). Saline and diluent controls (#1-ND 800, Eli Lilly LLC, IN, US) were used as blanks. Fresh insulin served as the negative control. Two series of IDF (-) and IDF (+) samples were prepared at the following dilution ratios: 1/10, 1/20, 1/40, and 1/100. For each well (2 µL of sample), 0.5 µL of the ThT working solution and 197.5 uL of the PBS were combined to achieve a final ThT concentration of 1 µM. Plates were gently mixed and incubated for 10–30 min at RT in the dark.

Fluorescence was then measured using a microplate reader (Synergy H1, BioTek Inc., VT, US) at excitation wavelengths of 440–450 nm and emission wavelengths of 480–490 nm.

Raw fluorescence values were corrected by subtracting the corresponding blank-control background. The percentage of fibril content was calculated according to the following equation:

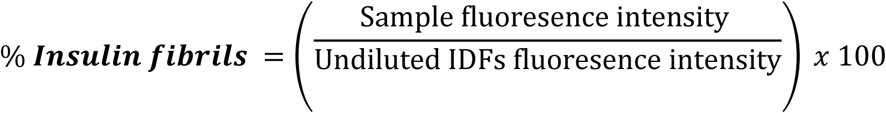

Each experiment was performed independently at least three times in triplicate.

### 2.4 Characterization of fibrils using a particle analyzer

To analyze fibrils size, 3 μL of IDF (+), IDF (-) or saline (0.9%) were loaded into the cartridge chamber. The following cartridges were employed: C400 for particles size ∼ 70 nm-400 nm (1 x 10^7^- 1 x 10^11^) and C900 for ∼ 130nm-900 nm (1 x 10^6^- 5 x 10^10^) to encompass all fibrils using Spectradyne’s nCS1™ particle analyzer (Spectradyne LLC, CA, US). Each sample was inserted as a single replicate and repeated three times from different fibril samples.

### 2.5 Differentiation of Bone Marrow-Derived Macrophages (BMDMs)

Bone marrow-derived macrophages were isolated and differentiated following established protocols ^24^. Briefly, mice were euthanized by CO₂ inhalation followed by cervical dislocation. Femurs were aseptically harvested and flushed with sterile phosphate-buffered saline (PBS) to collect bone marrow cells.

Cells were cultured in RPMI-1640 medium (#11835-030, Thermo Fisher Scientific) supplemented with 10% fetal bovine serum (FBS) (#16140-071, Thermo Fisher Scientific), 1% penicillin-streptomycin (#15140-122, Thermo Fisher Scientific), and 20% L929 cell-conditioned medium as a source of macrophage colony-stimulating factor (M-CSF).

Cultures were maintained at 37°C in a humidified atmosphere containing 5% CO₂ for 7 days without media changes to allow complete differentiation. On day 7, adherent macrophages were harvested and used immediately for studies.

### 2.6 Cytotoxicity assessment using MTT Assay

BMDMs were seeded in flat-bottom 96-well tissue culture plates (#FB012931, Thermo Fisher Scientific) at a density of 5 × 10⁴ cells/well in 200 μL of complete RPMI-1640 medium. After 24 hours of incubation, cells were treated with IDF (+), IDF (-), or m-cresol at serial dilutions (1/3 – 1/192). Media and serial dilutions of saline were used as untreated controls. Treatments were applied for 1 hour and 6 hours at 37°C in 5% CO₂. Following treatment, cell viability was assessed using the (3-[4,5-dimethylthiazol-2-yl]-2,5-diphenyl tetrazolium bromide (MTT) assay (#30-1010, ATCC, VA, US). Briefly, 15 μL of MTT reagent was added to each well (150 μL), and plates were incubated for 3 hours at 37°C in 5% CO₂, protected from light. After incubation, plates were centrifuged at 1,800 × g for 15 minutes at RT. Supernatants were carefully removed, and 200 μL of dimethyl sulfoxide (DMSO) (#D8418, Sigma-Aldrich) was added to each well. Plates were incubated for an additional 10 minutes at 37°C and then mixed. Absorbance was measured at 570 nm using a microplate reader (SpectraMax, Molecular Devices, CA, US).

Cell viability was calculated using the following formula:

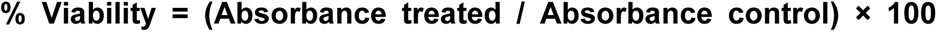. Each experiment was performed in triplicate wells and repeated at least three times.

### 2.7 Measurement of intracellular Reactive Oxygen Species (ROS)

Intracellular ROS levels were measured using the cellular ROS assay kit (#ab186027, Abcam, MA, US) according to the manufacturer’s protocol with minor modifications. BMDMs were seeded in black-walled, clear-bottom 96-well plates (#3603, Corning Costar®) at a density of 3.5 × 10⁴ cells per well. After 24 hours, cells were gently washed three times with pre-warmed RPMI-1640 medium supplemented with 10% FBS and 1% penicillin-streptomycin. Fresh supplemented medium (100 μL) was added to each well, followed by 100 μL of the ROS detection reagent. Plates were incubated for 50 minutes at 37°C in 5% CO₂. Subsequently, cells were treated with IDF (+), IDF (-) at 1/100, or m-cresol solution at 1/3 or 1/100 dilution. The medium served as the negative control, and hydrogen peroxide (H₂O₂) (1 mM) served as the positive control. Cells were incubated for an additional 30 minutes or 1 hour at 37°C in 5% CO₂. Fluorescence intensity was measured using a microplate reader (SpectraMax, Molecular Devices) in bottom-read mode with excitation/emission wavelengths of 520/605 nm. Background fluorescence was subtracted using wells containing only medium with reagent or medium with reagent plus treatments (no cells). Fluorescence data were analyzed as Relative Fluorescence Units (RFU) compared to untreated cells. Treatments were performed in triplicate wells and repeated in at least three independent experiments.

### 2.8 Assessment of Pro-Inflammatory Cytokine MIP-1α by ELISA

BMDM (∼5.5 × 10⁴ cells/well) plated in a 96-well tissue culture plate (#FB012931, Thermo Fisher Scientific) were exposed to IDF (+), IDF (-) at dilution 1/100, or m-cresol at dilution 1/3 for 6 or 24 hours. Following incubation, cell culture supernatants were collected and stored at −80°C until use. Levels of the pro-inflammatory cytokine MIP-1α in the supernatants were quantified using the Mouse MIP-1α DuoSet ELISA kit (#DY450-05, R&D Systems, MN, USA), according to the manufacturer’s instructions.

### 2.9 RNA Extraction and cDNA Synthesis

BMDMs were seeded in 6-well tissue culture plates (#3506, Corning Costar®) at a density of around 5 × 10⁵ cells/well. After 24 hours, cells were treated with IDF (+) or IDF (-) at 1/100 dilution, or m-cresol at 1/3 dilution for 1 hour. Following treatment, total RNA was extracted using TRIzol™ Reagent (#15596026, Thermo Fisher Scientific) according to the manufacturer’s protocol.

Briefly, culture medium was aspirated, and 1 mL of TRIzol™ Reagent was added directly to each well. Cells were lysed by pipetting up and down several times, and lysates were incubated at RT for 5 minutes to ensure complete dissociation of nucleoprotein complexes. Then, 200 μL of chloroform (#288306, Sigma-Aldrich) and 10 μL of GlycoBlue™ Coprecipitant (#AM9515, Thermo Fisher Scientific) were added.

The mixture was vigorously hand shaken for ∼ 20 seconds, incubated at RT for 5 minutes, and centrifuged at 12,000 × g for 5 minutes at 4°C. The aqueous phase (approximately 500 μL) was transferred to a new 1.5 mL microcentrifuge tube.

RNA was precipitated by adding 500 μL of isopropanol (#I9516, Sigma-Aldrich), gently mixed, and incubated at RT for 10 minutes. Samples were centrifuged at 12,000 × g for 10 minutes at 4°C to pellet the RNA. The supernatant was discarded, and the RNA pellet was washed with 1 mL of 75% ethanol (#BP2818, Thermo Fisher Scientific), vortexed briefly and centrifugated at 7,500 × g for 5 minutes at 4°C.

Ethanol was removed and the RNA pellet was air-dried at RT for approximately 10 minutes. The pellet was resuspended in 15–20 μL of RNase-free water and incubated at 60°C for 10 minutes to facilitate complete dissolution.

RNA concentration and purity were assessed using a NanoDrop 2000/2000c spectrophotometer (Thermo Fisher Scientific).

For cDNA synthesis, 1 μg of total RNA was reverse transcribed using the iScript™ cDNA Synthesis Kit (#1708891, Bio-Rad, CA, US) in a 20 μL reaction volume, according to the manufacturer’s instructions. The reverse transcription reaction was performed on a thermal cycler with the following program: 25°C for 5 minutes (priming), 46°C for 20 minutes (reverse transcription), and 95°C for 1 minute (inactivation). cDNA samples were stored at -20°C until utilized.

### 2.10 Quantitative Real-Time PCR (RT-qPCR)

Gene expression analysis was performed using SYBR Green-based quantitative real-time PCR. For each reaction, a 20 μL master mix was prepared containing: 10 μL Fast SYBR™ Green Master Mix (#4385612, Thermo Fisher Scientific), 1 μL forward primer (500 nM), 1 μL reverse primer (500 nM), 7 μL RNase-free water, 1 μL cDNA template. Primers were purchased pre-designed from the Integrated DNA Technologies company (IDT) (IA, US). Sequences for target genes are provided in Table S1 (*See supplementary material*). The master mix was dispensed into a MicroAmp® Fast Optical 96-well reaction plate (#4346907, Thermo Fisher Scientific), and reactions were carried out on a StepOnePlus™ Real-Time PCR System (Applied Biosystems, Thermo Fisher Scientific) using the following thermal cycling conditions: initial denaturation at 95°C for 20 seconds, followed by 40 cycles of denaturation at 95°C for 3 seconds and annealing/extension at 60°C for 30 seconds. Amplification data were analyzed using StepOne Software v2.3 (Applied Biosystems). Relative gene expression was calculated using the comparative Ct (2−ΔΔCt) method ^25^, with β-actin (ACTB) serving as the endogenous reference gene for normalization. The genes evaluated included inflammatory cytokines: TNF-α, IL-6, TGF-β; JAK/STAT pathway: JAK1, STAT1, STAT6; NF-κB pathway: IkBα; antioxidant response: KEAP1, NRF2. Each sample was analyzed in duplicate, and experiments were repeated with at least four independent biological replicates.

The housekeeping gene ACTB was selected as a control because it demonstrated stable Ct values within a consistent range of 14–17 across all samples and experiments, ensuring reliable and reproducible measurements as compared to GAPDH (data not shown) for the tested conditions.

### 2.11 Statistical Analysis

Data are depicted as mean ± standard error of the mean (SEM) or mean ± standard deviation (SD), as specified in the figure legends. Statistical analyses were conducted using GraphPad Prism version 11.0. For comparisons among multiple groups, one-way or two-way analysis of variance (ANOVA) followed by Tukey’s post hoc test for multiple comparisons was used. For comparisons between two groups, unpaired Student’s t-test was employed, as indicated in the figure legends. Statistical significance was considered at: *p < 0.05, **p < 0.01, ***p < 0.001, ****p < 0.0001. All experiments included at least three independent biological replicates in three independent experiments.

## 3 Results

### 3.1 Quantification of Insulin fibrils by ThT assay

To determine whether the presence of preservatives in the insulin diluent alters fibril abundance or interferes with ThT-based detection, IDF (+) and IDF (-) preparations were quantified across a series of serial dilutions (Figure 1).

**Figure 1:**
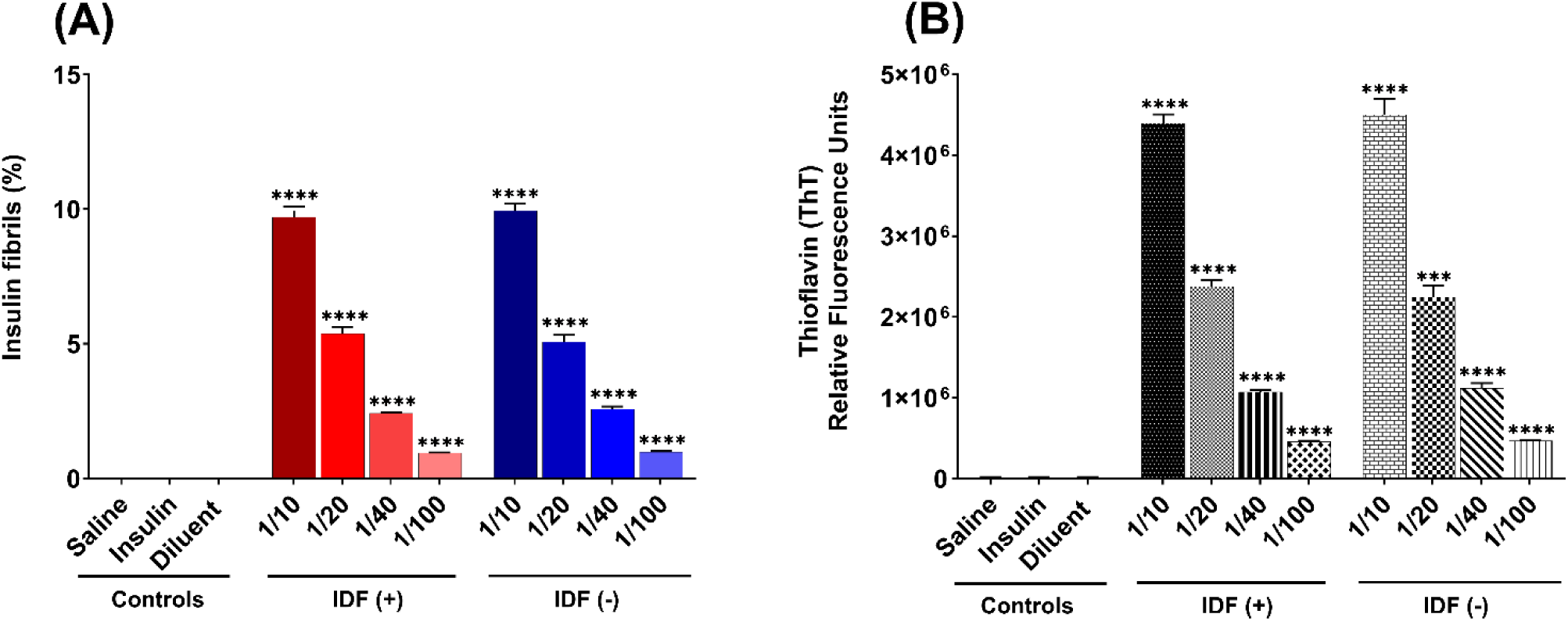
Quantification of insulin-derived fibrils (IDF(+) and IDF(-)) by thioflavin T (ThT) fluorescence, demonstrating that phenolic preservatives neither alter fibril content nor interfere with ThT binding or fluorescence output. IDF(+) and IDF(-) show **(A)** similar percentages of fibril content and **(B)** similar relative fluorescence intensity. The data are presented as the mean ± SEM of three independent experiments. ***p < 0.001, ****p < 0.0001 as determined by unpaired t-test.

Normalization of ThT fluorescence to the signal from undiluted (100%) IDFs demonstrated that the calculated fibril percentages closely matched the expected dilution ratios in both fibril groups. This proportional decline indicates that ThT reliably captures fibril abundance across a broad concentration range (Figure 1A). Notably, direct comparison of IDF (+) and IDF (-) samples at equivalent dilution ratios revealed comparable fibril percentages, indicating that preservatives in the IDF preparation do not affect the ThT response and that both fibril types induce similar levels of protein aggregation.

The fluorescence data demonstrated the same dilution-dependent pattern and group similarities as the normalized fibril quantification, thus confirming that preservatives did not affect ThT signal generation (Figure 1B), which is consistent with the normalized results in Figure 1A. Controls demonstrated fluorescence near zero, indicating that neither matrix produced false-positive ThT signals. In contrast, both IDF samples exhibited fluorescence intensities well above that of the controls, confirming the presence of ThT-reactive fibrillar structures. The strong concordance between normalized fibril percentages and raw fluorescence traces demonstrates that preservatives have no significant effect on fibril content and do not interfere with ThT binding or fluorescence output.

### 3.2 IDF with and without preservative presence have similar particle sizes

To determine whether saline, preservative-containing insulin, or IDF formulations contribute to detectable particulate species, we quantified particle-size distributions and total particle concentrations (Figure 2).

**Figure 2:**
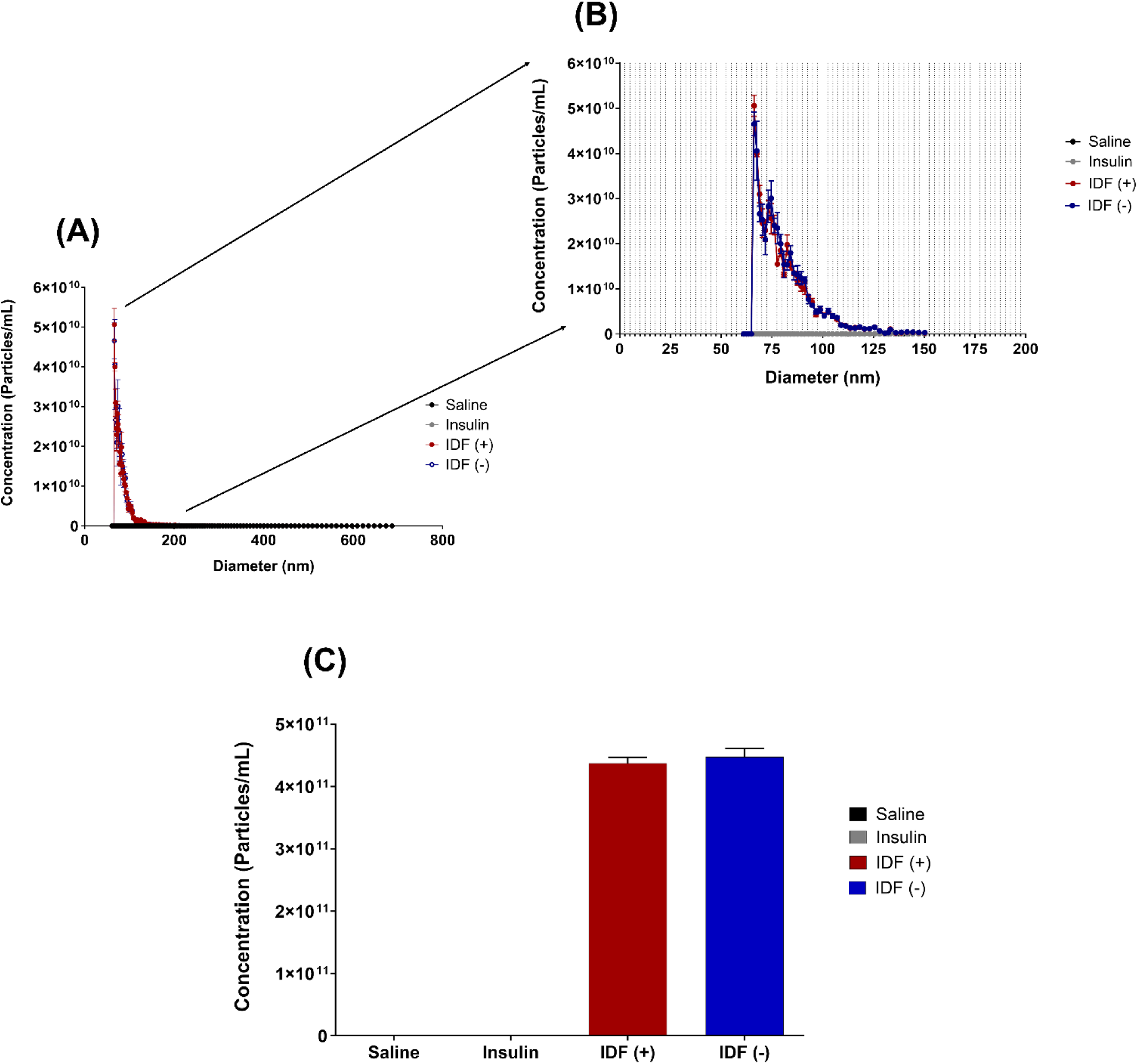
Particle size analysis of insulin-derived fibrils (IDF(+) and IDF(-)) by microfluidic profiling, demonstrating that phenolic preservatives neither interfere with fibril detection nor affect particle size. IDF(+) and IDF(-) show **(A)** similar particle size distributions **(B)** around 75-100 nm and **(C)** similar total particle concentrations. Data are presented as mean ± SEM of three independent experiments. Statistical significance was determined by one-way ANOVA with Tukey’s as a post-test for multiple comparisons.

Across the full diameter spectrum measured, saline and insulin exhibited nearly identical particle-size distribution curves (Figure 2A). Both samples remained at baseline across all measured diameters, and no enrichment of large-diameter particles (>100 nm) was observed in the insulin group. These findings demonstrate that native insulin preparations do not spontaneously generate fibrils or high-molecular-weight aggregates under the tested conditions. Moreover, these data demonstrate that the preservatives included in commercial insulin formulations do not produce detectable particulate artifacts, thus excluding false-positive signals attributable to formulation excipients.

In contrast, both IDF (+) and IDF (-) samples displayed particle counts several orders of magnitude higher than saline and insulin, although superimposable between both fibrils, with a pronounced enrichment of larger-diameter species characteristic of insulin fibrils (Figure 2A). Consistently, total particle concentrations in IDF samples were markedly elevated relative to both controls (p < 0.0001 for IDF (+) and p < 0.01 for IDF (-)) (Figure 2B), thus confirming that IDF preparations contain abundant fibrillar structures.

The absence of detectable differences between IDF groups indicates that preservatives neither alter fibril abundance nor affect the assay’s ability to quantify fibrillar particles. This observation provides strong evidence that preservatives do not interfere with fibril detection nor contribute measurable particulate background in the microfluidic analysis and that the preservative may not impact the size of fibril formation.

### 3.3 IDF (+) shows increased cytotoxicity

To evaluate fibril cytotoxicity, we exposed BMDMs to IDF (+), IDF (-), or m-cresol solution at serial dilutions for 1 or 6 hours. Cell viability was assessed using the MTT assay (Figure 3). After 1 hour, m-cresol at a 1/3 dilution demonstrated significant cytotoxicity, reducing cell viability to approximately 18% compared with the untreated media control (Figure 3A). In contrast, both IDF (+) and IDF (-) displayed modest cytotoxicity at this early stage, with no significant difference between the two fibril types at the same concentration. However, after 6 hours, IDF (+) induced significantly greater cytotoxicity (∼30% viability) than IDF (-) (∼45% viability) at the same concentration (Figure 3B). After 24 h, IDF (+) still exhibited greater cytotoxicity (12.45% viability) than IDF (−) (33.04% viability) at equivalent dilution (*See supplementary Figure S1*). The time-dependent increase in toxicity implies that phenolic preservatives within IDF (+) enhance macrophage cytotoxicity during prolonged exposure, while both fibril preparations exhibit similar toxicity at lower dilutions. Collectively, these results demonstrate that IDF possess inherent cytotoxic activity toward macrophages and that phenolic preservatives such as m-cresol can potentiate this effect markedly over time.

**Figure 3:**
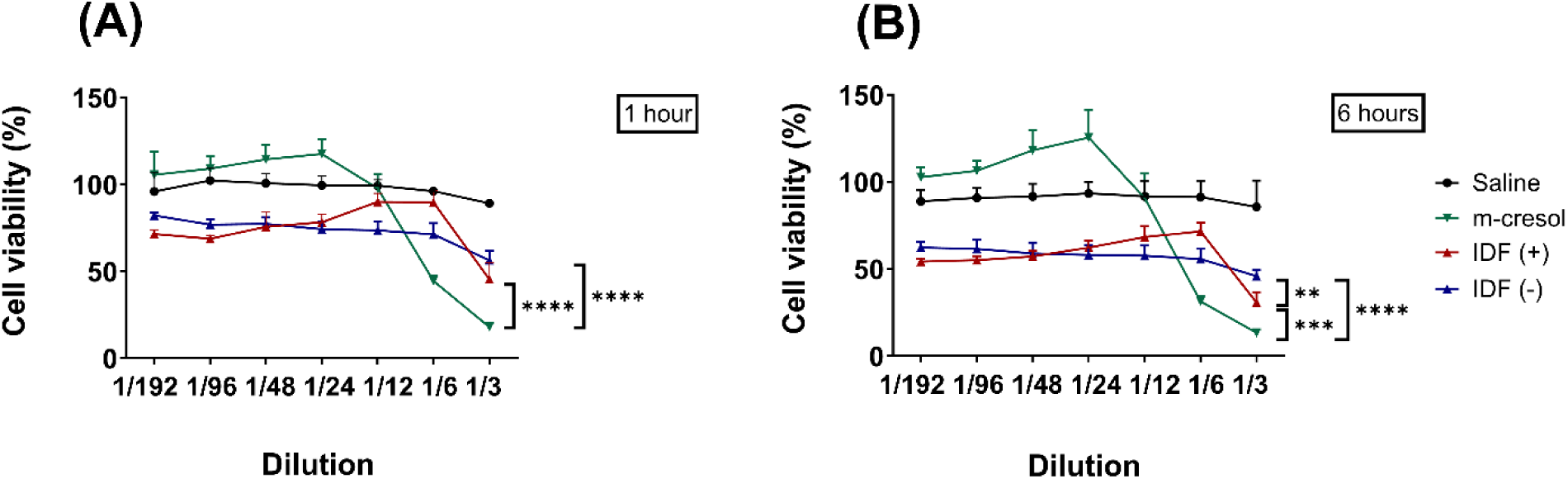
Cytotoxic effects of insulin-derived fibrils (IDFs) and m-cresol in BMDMs as a function of exposure time and concentration, with IDF(+) exhibiting increased cytotoxicity relative to IDF(-) at higher dilutions. Bone Marrow-Derived Macrophages (BMDMs) were exposed to IDF (+), IDF (-), or m-cresol solution (3.15 mg/mL) at serial dilutions for **(A)** 1 hour and **(B)** 6 hours. Cell viability was measured using the MTT assay (3-[4,5-dimethylthiazol-2-yl]-2,5-diphenyl-tetrazolium bromide). Results are shown as a percentage of untreated controls. Data are presented as the mean ± SD of three independent experiments performed in triplicate. **p < 0.01, ***p < 0.001, ****p < 0.0001, as determined by two-way ANOVA with Tukey’s post hoc test for multiple comparisons.

### 3.4 Fibrils formed in the presence of phenolic preservatives induce ROS in macrophages

ROS play diverse roles in macrophage biology, acting as antimicrobial agents, signaling molecules, and mediators of oxidative stress ^26,27^. To assess whether IDF stimulate ROS production, we measured intracellular ROS levels in BMDMs after treatment with IDF (+), IDF (-), or m-cresol solution (Figure 4).

**Figure 4.**
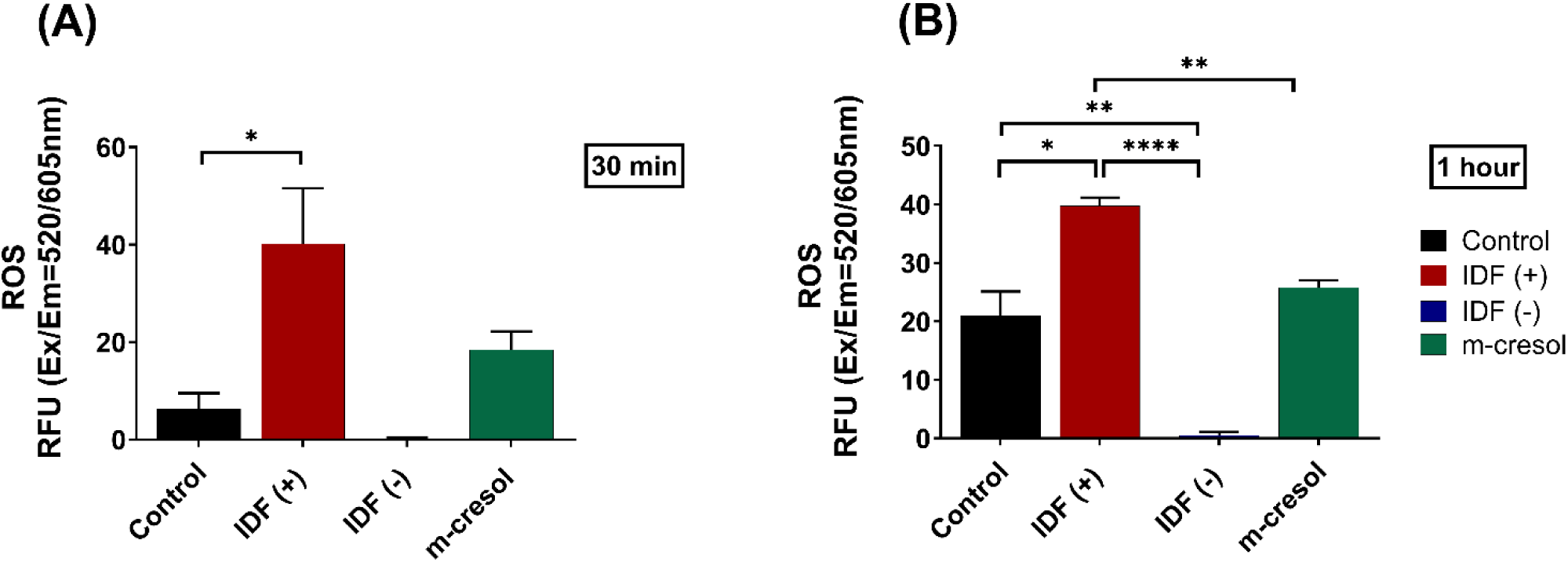
Intracellular reactive oxygen species (ROS) production in Bone Marrow-Derived Macrophages (BMDMs) following stimulation with insulin-derived fibrils (IDFs) and m-cresol. IDF(+) induces significantly higher ROS levels than IDF(-) and m-cresol. BMDMs were treated with IDF (+), IDF (-), or m-cresol at a 1/100 dilution for **(A)** 30 minutes and **(B)** 1 hour. Intracellular ROS levels were measured using a fluorescence-based assay. Data are presented as the mean ± SEM of three independent experiments performed in triplicate. *p < 0.05, **p < 0.01, ****p < 0.0001 as determined by unpaired t-test.

After 30 minutes of exposure at a 1/100 dilution, IDF (+) significantly increased intracellular ROS levels compared to control (Figure 4A). In contrast, IDF (-) and m-cresol did not cause a significant increase in ROS at this time point.

At 1-hour post-treatment, IDF (+) continued to induce strong ROS production, significantly exceeding that triggered by both IDF (-) and the m-cresol solution (Figure 4B). Interestingly, m-cresol at a lower dilution (1/3) (*See supplementary Figure S2*) induced ROS levels comparable to those caused by IDF (+) at 1/100 dilution, indicating that elevated preservative levels can provoke oxidative stress independently.

These findings demonstrate that fibrils formed in the presence of phenolic preservatives enhance macrophage activation by promoting ROS generation. They also suggest a synergistic interaction between fibrils and preservatives, as lower concentrations of IDF (+) are sufficient to induce significant ROS production. In contrast, m-cresol alone requires much higher concentrations to achieve similar effects. The minimal ROS response to IDF (-) indicates that fibrils alone do not trigger ROS strongly, and that their interaction with preservatives likely amplifies oxidative stress.

### 3.5 Exposure to IDF induces increased MIP-1a levels

To assess the inflammatory potential of IDFs, we measured MIP-1α (CCL3) levels in BMDM supernatants. Both IDF (+) and IDF (-) induced increased MIP-1α production compared to the control after 6 hours, with no significant difference between the two fibril conditions (Figure 5A). After 24 hours, only IDF (+) maintained significantly elevated MIP-1α levels relative to the control (Figure 5B).

**Figure 5:**
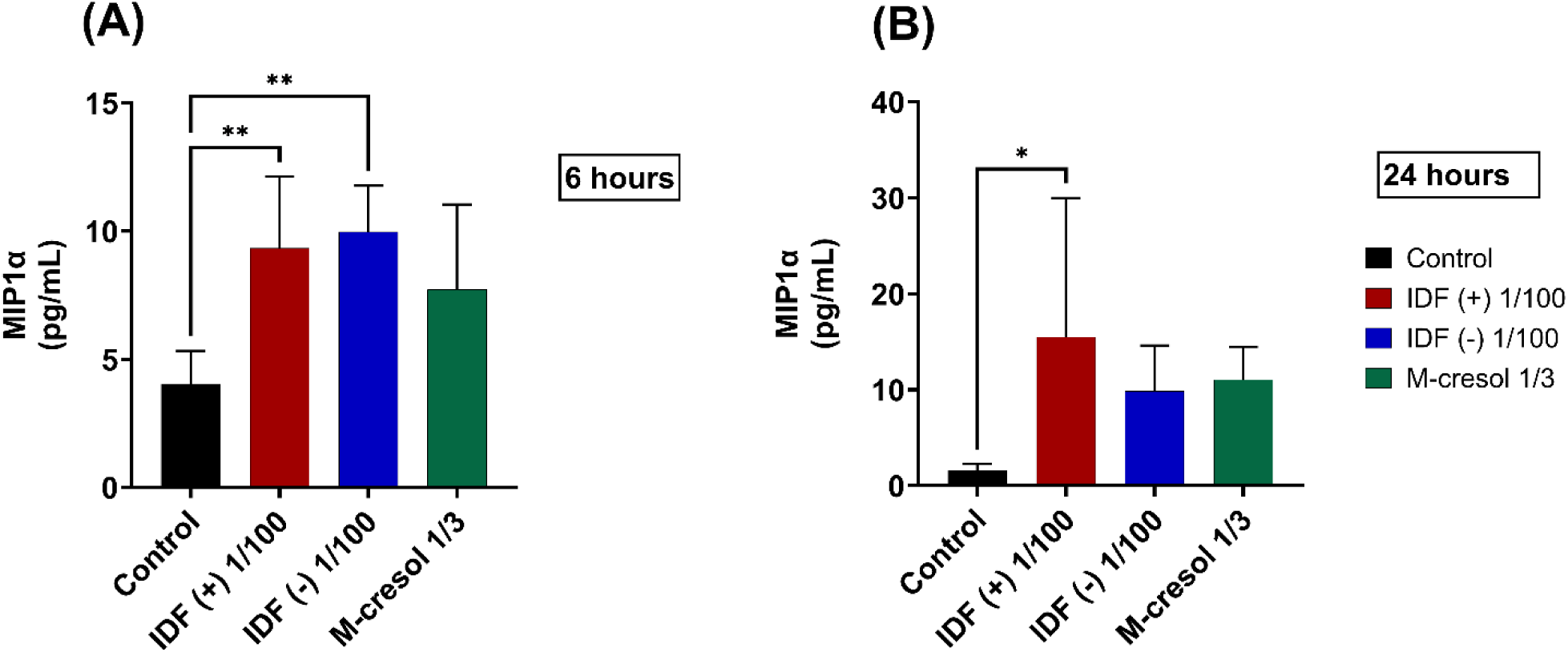
IDF (+) and IDF (-) induce MIP-1α release in Bone Marrow Derived Macrophages (BMDM). MIP-1α chemokine expression in IDF (+) and IDF (-) after **(A)** 6 hours and **(B)** 24 hours. ELISA assay was performed in duplicate of at least six independent experiments. Data are presented as mean ± SD. * p< 0.05 **p < 0.01 as determined by one-way ANOVA with Tukey’s for multiple comparisons.

We also detected mRNA levels of key pro-inflammatory cytokines, TNF-α, and the immunoregulatory cytokine TGF-β in BMDMs after 1 hour of treatment (*See supplementary Figure S3*). Only IDF (-), increased TNF-α expression as compared to untreated control (dashed line) (Figure S3A). whereas m-cresol downregulated TGF-β compared to control and IDF (+) (Figure S3B).

Together, these findings indicate that IDFs activate macrophages towards distinct pro-inflammatory profiles. Furthermore, they suggest that even low concentrations of insulin fibrils can trigger inflammatory responses in macrophages. The downregulation of TGF-β by m-cresol further reinforces distinct immunomodulatory mechanisms.

### 3.6 Fibrils modulate distinct signaling pathways, suggesting ROS-independent mechanisms

We further investigated whether both fibrils modulate key inflammatory signaling pathways in macrophages differently (Figure 6). After 1 hour of treatment, only IDF (-) increased NRF2 mRNA expression significantly as compared to media-only control (p< 0.01) (Figure 6A). M-cresol significantly reduced STAT6 mRNA expression compared to control (p< 0.05), which remained below baseline (Figure 6B). The contrasting gene expression pattern implies that the fibrils activate pathway components not solely driven by ROS, whereas the presence of m-cresol appears to suppress components of this response.

**Figure 6.**
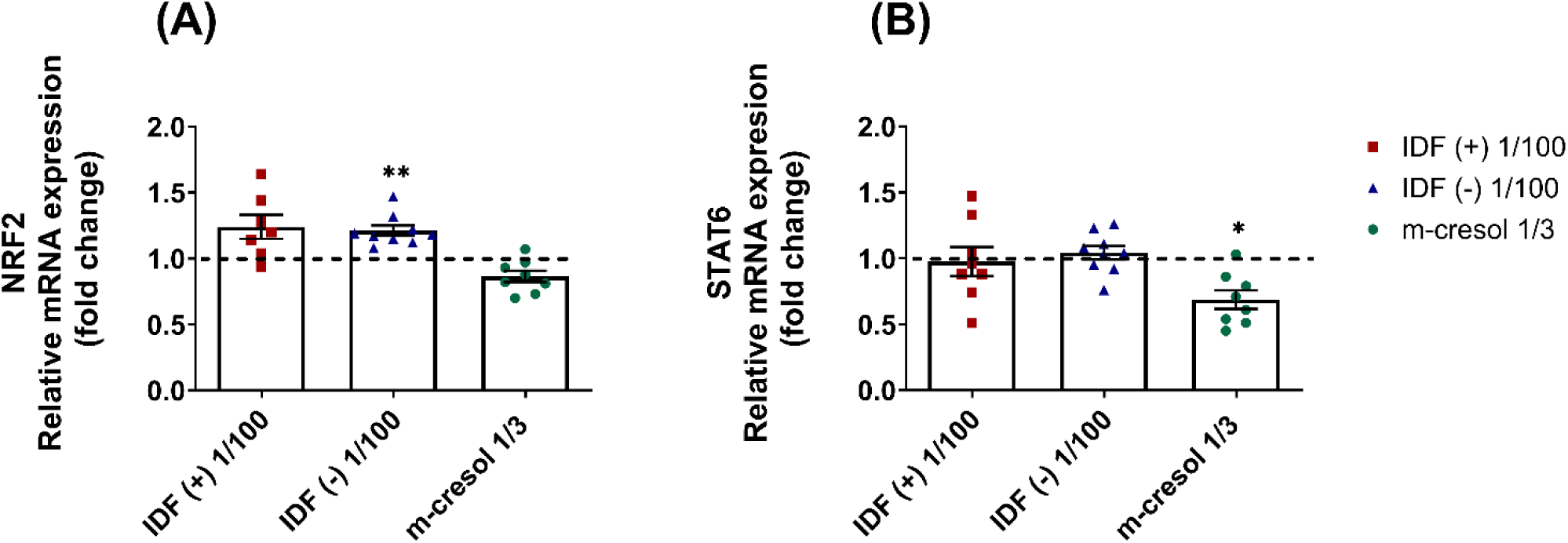
Inflammatory signaling pathways modulated in bone marrow–derived macrophages (BMDMs) following stimulation with insulin-derived fibrils (IDFs) and m-cresol. Relative mRNA expression levels of **(A)** NRF2 and **(B)** STAT6 were quantified by RT-qPCR. Data are presented as mean ± SEM of fold change relative to untreated control (control = 1, shown as dashed line) and represent at least four independent experiments performed in duplicate. Statistical analysis was performed using ΔCt values, and the data was plotted as fold change. * p<0.05, **p < 0.01, as determined by Mixed-effect analysis or One-way ANOVA with Tukey’s post hoc test for multiple comparisons.

## 4 Discussion

Although insulin fibrils have been associated with insulin infusion set complications and local inflammation ^5,14^, the mechanisms by which IDF trigger immune activation remain incompletely elucidated. While phenolic preservatives are well recognized for their cytotoxic potential ^28,29^, these data demonstrate that IDFs themselves exert pronounced cytotoxic and immunomodulatory effects on BMDMs. Specifically, fibrils reduced macrophage viability in a concentration- and time-dependent manner, with fibrils formed in the presence of phenolic preservatives (IDF(+)) displaying significantly greater cytotoxicity than those formed in their absence (IDF(-)).

### 4.1 Oxidative Stress and ROS Generation

Amyloid fibrils are known inducers of reactive ROS ^30,31^, which function as key regulators of cellular signaling, microbial clearance, differentiation, and gene expression ^32–34^. Notably, IDF(+) induced substantially higher levels of ROS in macrophages when compared with both IDF(-) and m-cresol at equivalent dilutions. At higher concentrations, m-cresol alone also induced significant ROS, implicating a potential synergistic interaction between phenolic preservatives and insulin fibrils in promoting oxidative stress. This enhanced ROS generation may contribute to local inflammatory responses at infusion sites.

### 4.2 Fibril-Induced Upregulation of MIP-1α

MIP-1α is secreted by activated macrophages and plays an important role in leukocyte recruitment during tissue injury and inflammation ^35^. In particular, this chemokine is predominantly released by M1 macrophages and is associated with a pro-inflammatory phenotype ^36^. The increased release of MIP-1α by IDFs further supports their inflammatory effect and suggests potential mechanisms through which fibrils promote inflammation. Moreover, the sustained release of MIP-1α over 24 h in the IDF (+) group indicates a persistent inflammatory response, which is possibly synergistic with m-cresol.

### 4.3 Signaling Pathways

We assessed whether differential ROS generation translated into distinct activation patterns. It is noteworthy that only IDF (-) positively modulated NRF2 (NF-E2-related factor-2) compared to the control. This transcription factor is associated with the regulation of the oxidative stress response, which we did not observe in IDF (-) in this study, and with anti-inflammatory effects ^37^. However, in macrophages, knocking down NRF2 showed mixed effects on inflammation and glucose metabolism ^38^. NRF2 has also been associated with inflammasome activation *in vivo* ^39^. Supporting this, IDF (-) also modulated the gene expression of TNF-α cytokine. Therefore, we speculate that NRF2 has an inflammatory role in this context.

Furthermore, m-cresol also significantly downregulated STAT6, which serves a role in the anti-inflammatory response by inducing a M2 type macrophages profile. Its inhibition is associated with increased acute inflammation ^40^. STAT6 has also been associated with TGF-β production^41^. Concordantly, TGF-β was downregulated in m-cresol compared to control and IDF (+). These results demonstrate that distinct fibrils and m-cresol can induce inflammation via distinct signaling pathways.

It should be noted that our analysis of signaling pathways was limited to gene-level expression data. Consequently, additional investigations at the protein level are necessary to confirm the biological relevance of these findings.

### 4.4 Mechanisms of Macrophage Activation

In summary, these findings suggest that macrophage activation by IDF is more likely to be mediated through direct receptor engagement, pattern-recognition signaling, or membrane perturbation rather than by oxidative stress alone. Phenolic preservatives may exacerbate inflammation indirectly through ROS-dependent cytotoxic stress or by modulating cell death pathways.

### 4.5 Role of Phenolic Preservatives in Fibril Bioactivity

A noteworthy finding is that phenolic preservatives did not alter fibril size distribution but did influence its biological activity. Preservatives, due to their amphipathic properties and known interactions with lipids and proteins ^42,43^, may modify fibril stability, surface hydrophobicity, or protein corona formation in ways that enhance cellular recognition and uptake. However, an alternative explanation that the observed effects are driven by m-cresol inducing inflammatory cell death resulting from membrane permeabilization and the release of damage- associated molecular patters (DAMPs) rather than immunomodulation cannot be excluded. Phenolic preservatives activate stress kinases and induce pro-inflammatory response through MCP-1 release ^44^. They are known to damage cell membrane, increase ROS generation, lead to protein denaturation, DNA and cytoskeleton damage and ferroptosis cell death ^45^ a type of cell death that can release DAMPs ^46^.

Elucidating the distinction between fibril formation (a structural process) and fibril bioactivity (a functional consequence) may be particularly relevant to understanding preservative-dependent inflammatory responses at infusion sites. Future studies employing advanced biophysical techniques, such as surface plasmon resonance, circular dichroism, or atomic force microscopy, may elucidate the mechanisms by which preservatives affect fibril response.

### 4.6 Clinical Implications

These results highlight the importance of fibril structure-activity relationships in driving immune activation and emphasize the need for formulation strategies that minimize fibril formation while preserving insulin biocompatibility. Minimal amounts of insulin fibrils may activate tissue-resident macrophages, thus initiating inflammatory cascades that could contribute to the formation of anti-insulin antibodies, lipohypertrophy, and other immune-mediated complications observed clinically. The preservative-dependent enhancement of fibril immunogenicity implies that novel preservative-free formulations or alternative stabilization strategies may reduce infusion-site inflammation and extend the lifespan of infusion sets.

### 4.7 Limitations and Future Directions

This study has several limitations that should be addressed in future investigations. First, this study utilized isolated murine BMDMs, which do not fully recapitulate the complexity of the tissue microenvironment or interactions with other immune cells such as T cells, dendritic cells, and fibroblasts. Studies using human monocyte-derived macrophages and tissue-resident macrophage models would be more clinically relevant. Second, while this fibrillation protocol was robust and reproducible, additional biophysical characterization, including secondary-structure analysis, fibril-length distribution, and surface-charge measurements, would strengthen mechanistic interpretations. Third, the exposures examined were relatively short-term (1–6 h), whereas chronic or repeated exposures may be a better approximation of the physiological conditions at infusion sites during prolonged pump use. Fourth, this investigation did not identify the specific receptors mediating fibril recognition or validate the molecular pathways through targeted genetic or pharmacological approaches. Candidate pathways such as Toll-like receptors, scavenger receptors (CD36, SR-A), and inflammasome components (NLRP3) warrant systematic investigation. Finally, *in vivo* models of subcutaneous insulin fibril exposure will be essential to validate these findings and assess the contribution of fibrils to those clinically observed infusion-site complications.

## 5 Conclusion

In summary, this study demonstrated that IDF induces significant cytotoxic and immunomodulatory effects in macrophages. Moreover, phenolic preservatives amplify these responses substantially through mechanisms that involve increased cytotoxicity and ROS generation. These findings provide mechanistic insight into inflammatory complications associated with insulin pump therapy. They suggest that formulation optimization, particularly regarding preservative and IDF selection or removal, is a viable strategy to improve the biocompatibility and longevity of insulin infusion systems.

## Acknowledgements

The authors thank Li Mao for performing the differentiation of bone marrow–derived macrophages (BMDMs) and supporting with preparing reagents for the experiment.

## Author Contributions

**P.S.C:** Conceptualization, Formal analysis, Methodology, Writing.

**C.C**: Methodology, Formal analysis, Writing.

**D.L.M**: Writing, Editing

**E.Z:** Formal analysis

**U.K:** Conceptualization, Original draft, Writing and editing, Funding acquisition.

## Conflicts of Interest

Ulrike Klueh is the founder and owner of Alva Innovations, Inc., which holds patent applications for methods of removing insulin fibrils from infusion paths. The research presented in this manuscript identifies problems that these patented technologies may address. This relationship did not influence the study design, data collection, analysis, or interpretation of the results. The other authors declare no conflicts of interest.

## Funding

This study was supported by the NIDDK Institute within the National Institute of Health (NIH) [grant number R01DK129681] and Digestive and Kidney Diseases (NIDDK) at the NIH [grant number 1R01DK133789].

## Statement AI use

During the preparation of this work, AI was used after the first draft to support language editing and improvements to sentence structure and manuscript flow. Subsequently, the manuscript was edited by the authors with expertise in writing scientific publications, such that the final version was not AI-generated. Furthermore, all content was reviewed, revised, and approved by all the authors, who accept full responsibility for the final version.

## Data Statement

The authors declare that all data supporting the findings of this study are provided within the manuscript and the supplementary materials.

## Glossary

CSII: Continuous Subcutaneous Insulin Infusion
FDA: Food and Drug Administration
IDFs: Insulin-Derived Fibrils
BMDMs: Bone Marrow-Derived Macrophages
ROS: Reactive Oxygen Species
MIP-1α/CCL3: Macrophage Inflammatory Protein-1α
TNF-α: Tumor Necrosis Factor alpha
TGF-β: Transforming Growth Factor beta
STAT6: Signal Transducer and Activator of Transcription 6
NRF2: Nuclear Factor Erythroid 2-Related Factor 2
IIS: Insulin Infusion Set

