## Supplemental material for "Immunomodulatory Effects of Insulin-Derived Fibrils from Infusion Pumps: Role of Phenolic Preservatives in Macrophage Activation"

### Supplementary Material

| Gene | Forward Primer (5' → 3') | Reverse Primer (5' → 3') |
|---|---|---|
| **TNF-α** | TCTGTCTACTGAACTTCGGGGTGATCG | GTATGAGATAGCAAATCGGCTGACGGTG |
| **IL-6** | CTTCCATCCAGTTGCCTTCTTG | AATTAAGCCTCCGACTTGTGAAG |
| **TGF-β** | GGAGAGCCCTGGATACCAAC | CAACCCAGGTCCTTCCTAAA |
| **JAK1** | GAACCACCTCAAGAAGCAGAT | GGCTTTCTTAGTGGCTACGAG |
| **STAT1** | TTGACAAAGACCACGCCTT | GACTTCAGACACAGAAATCAACTC |
| **STAT6** | GCCACCATCAGACAAATACTTC | AGTTCTTCCTGCTTCCGATG |
| **NFKBIA** | TGCCTGGCCAGTGTAGCAGTCTT | CAAAGTCACCAAGTGCTCCACGAT |
| **KEAP1** | GGAGTATATCTACATGCACTTCGG | GCAGCGTACGTTCAGATCA |
| **NFE2L2** | TGATGGACTTGGAGTTGCC | TCAAACACTTCTCGACTTACTCC |
| **ACTB (β-actin)** | GATTACTGCTCTGGCTCCTAG | GACTCATCGTACTCCTGCTTG |

**Table S1.** Primer sequences used for RT-qPCR analysis

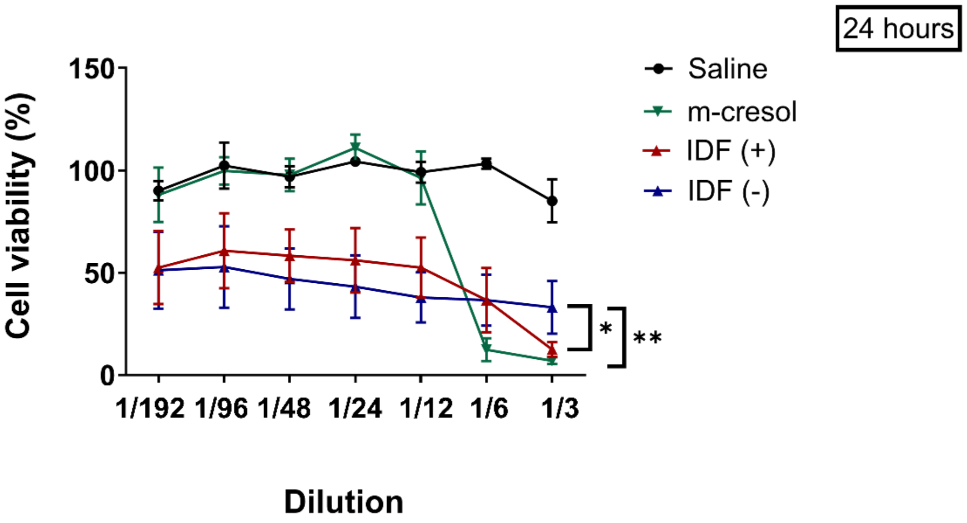


**Figure S1.** Cytotoxic effects of insulin-derived fibrils (IDFs) and m-cresol in BMDMs as a function of exposure time and concentration, with IDF(+) exhibiting increased cytotoxicity relative to IDF(-) at higher dilution. Bone Marrow-Derived Macrophages (BMDMs) were exposed to IDF (+), IDF (-), or m-cresol solution (3.15 mg/mL) at serial dilutions for 24h. Cell viability was measured using the MTT assay (3-[4,5-dimethylthiazol-2-yl]-2,5-diphenyl-tetrazolium bromide). Results are depicted as a percentage of untreated controls. Data represent the mean ± SD of at least two independent experiments performed in triplicate. $^{**}p < 0.01$, $^{***}p < 0.001$, $^{****}p < 0.0001$, as determined by two-way ANOVA with Tukey's post hoc test for multiple comparisons.

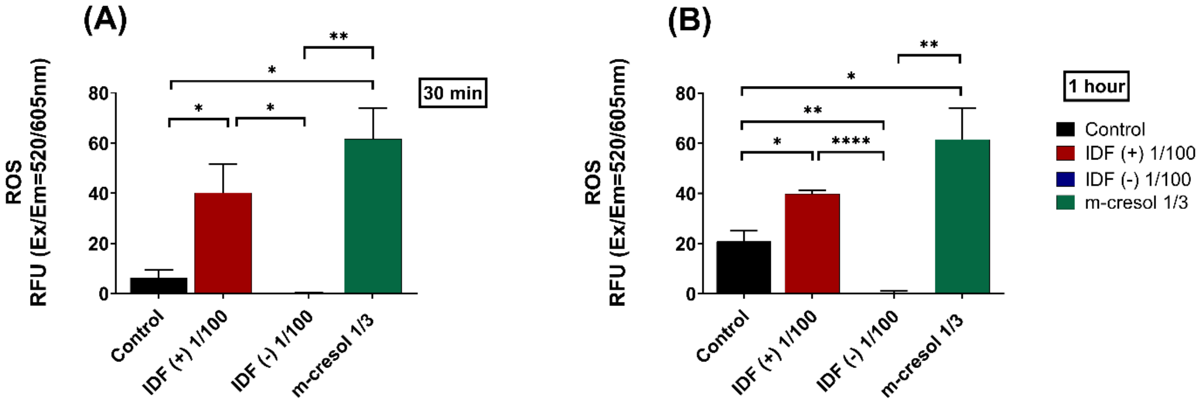


**Figure S2.** Intracellular reactive oxygen species (ROS) production in Bone Marrow–Derived Macrophages (BMDMs) following stimulation with insulin-derived fibrils (IDFs) and m-cresol. IDF (+) at dilution 1/100 induced significantly higher ROS levels than IDF (−) and produced ROS levels comparable to m-cresol at 1/3 dilution. BMDMs were treated with IDF (+), IDF (-), or m-cresol for **(A)** 30 minutes and **(B)** 1 hour. Intracellular ROS levels were measured using a fluorescence-based assay. Data represent mean ± SEM of three independent experiments performed in triplicate. *p < 0.05, **p < 0.01, ****p < 0.0001 as determined by unpaired *t*-test.

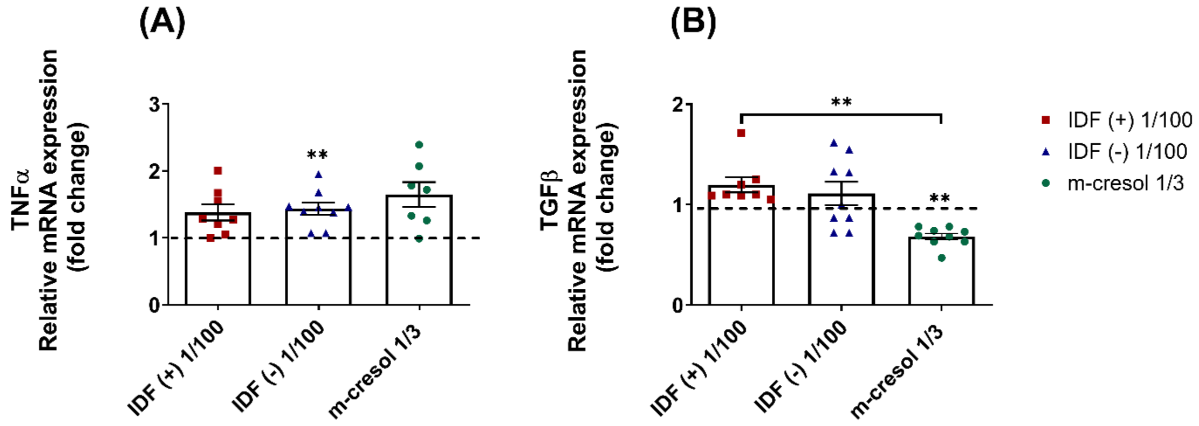


**Figure S3.** Cytokine gene expression profiles in bone marrow–derived macrophages (BMDMs) following stimulation with insulin-derived fibrils (IDFs) and m-cresol. IDF(-) significantly induces TNF-α whereas m-cresol significantly reduced TGF-β compared to control and IDF(+) 1/100. BMDMs were treated with IDF (+), IDF (-), or m-cresol for 1 hour. Relative mRNA expression levels of **(A)** TNFα, and **(B)** TGFβ were quantified by RT-qPCR. Data are presented as mean ± SEM of fold change relative to the untreated control (control = 1, shown as a black dashed line) from at least four independent experiments performed in duplicate. Statistical analysis was performed using ΔCt values, and the data was plotted as fold change. **p < 0.01, as determined by Mixed-effect analysis or One-way ANOVA with Tukey's post hoc test for multiple comparisons.
